# Chromosome-level genome assembly of the hydrozoan jellyfish *Cladonema pacificum*

**DOI:** 10.64898/2026.11.26.769559

**Authors:** Junichiro Ui, Katsushi Yamaguchi, Hiroko Nakatani, Koto Kon-Nanjo, Tetsuo Kon, Yuu Ishii, Noriyo Takeda, Ryusaku Deguchi, Shuji Shigenobu, Yu-ichiro Nakajima

## Abstract

The hydrozoan jellyfish *Cladonema pacificum*, which possesses experimentally tractable polyp and medusa stages, is an emerging cnidarian model for studies of regeneration, development, physiology, and behavior. Although experimental techniques have been established for this species, the lack of a chromosome-level reference genome has limited genome-wide and comparative analyses. Here, we present the first chromosome-level genome assembly of *C. pacificum* generated using HiFi long-read sequencing and Hi-C technology. The representative haploid assembly comprises 15 chromosome-scale scaffolds and 121 additional scaffolds, spanning 504.0 Mb with a scaffold N50 of 32.1 Mb. Gene annotation integrating RNA-seq, PacBio Iso-Seq, and protein homology identified 17,206 protein-coding genes, of which 95.7% received at least one functional annotation. Phylogenomic and macrosynteny analyses further confirmed the placement of *C. pacificum* within Hydrozoa and revealed extensive chromosome-scale conservation with other hydrozoan genomes. This genomic resource provides a valuable foundation for comparative genomics and future molecular, cellular, and evolutionary studies using *C. pacificum* as an emerging hydrozoan model.

## Background & Summary

Cnidarians are the sister group to bilaterians and occupy an important phylogenetic position for understanding the early evolution of animal body plans, developmental programs, and life cycles^1,2^. In addition to their remarkable regenerative capacities, many cnidarian lineages, particularly jellyfish, exhibit complex life cycles comprising planula larvae, polyps, and medusae. Over the past decade, genomic resources and experimental tools have expanded substantially for a limited number of cnidarian model organisms, including *Nematostella vectensis*^3,4^, *Hydra vulgaris*^5,6^, and *Hydractinia symbiolongicarpus*^7,8^. However, these established models are polyp-form cnidarians that lack a medusa stage. *Clytia hemisphaerica*, a hydrozoan jellyfish with a medusa stage, is also an established model in which genetic manipulation methods have been developed^9,10^, but its chromosome-level reference genome is not yet available. Although these model organisms have provided important insights into cnidarian biology, they represent only a limited portion of cnidarian diversity, particularly among medusa-bearing hydrozoans.

The hydrozoan jellyfish *Cladonema pacificum* is a promising emerging model in cnidarian research^11,12^. Its life cycle comprises a sexually reproducing medusa stage (Fig. 1A) and an asexually reproducing polyp stage (Fig. 1B). Gametes released by medusae develop through planula larvae into polyp colonies connected by stolons (Fig. 1C), from which medusae are asexually produced. Because the complete life cycle can be maintained under laboratory conditions^13,14^, mature medusae are available throughout the year for experimental studies. In addition, polyp colonies can enter a dormancy state at low temperatures, allowing long-term maintenance and subsequent recovery^14,15^. Furthermore, experimental techniques, including fluorescence *in situ* hybridization^16^ and electroporation-based siRNA-mediated knockdown^17^, enable both gene-expression and functional analyses in this species. *C. pacificum* has been used to investigate mechanisms of regeneration^18–21^, light-induced reproductive signaling^13,22,23^, tentacle branching morphogenesis^14,24^, and feeding behavior^25–27^. Together, these studies highlight *C. pacificum* as a versatile hydrozoan jellyfish model for investigating diverse biological processes. Despite its increasing utility as an experimental organism, a chromosome-level reference genome has not previously been available, thereby hindering genome-wide analyses of gene organization, regulatory architecture, and chromosome evolution.

**Figure 1.**
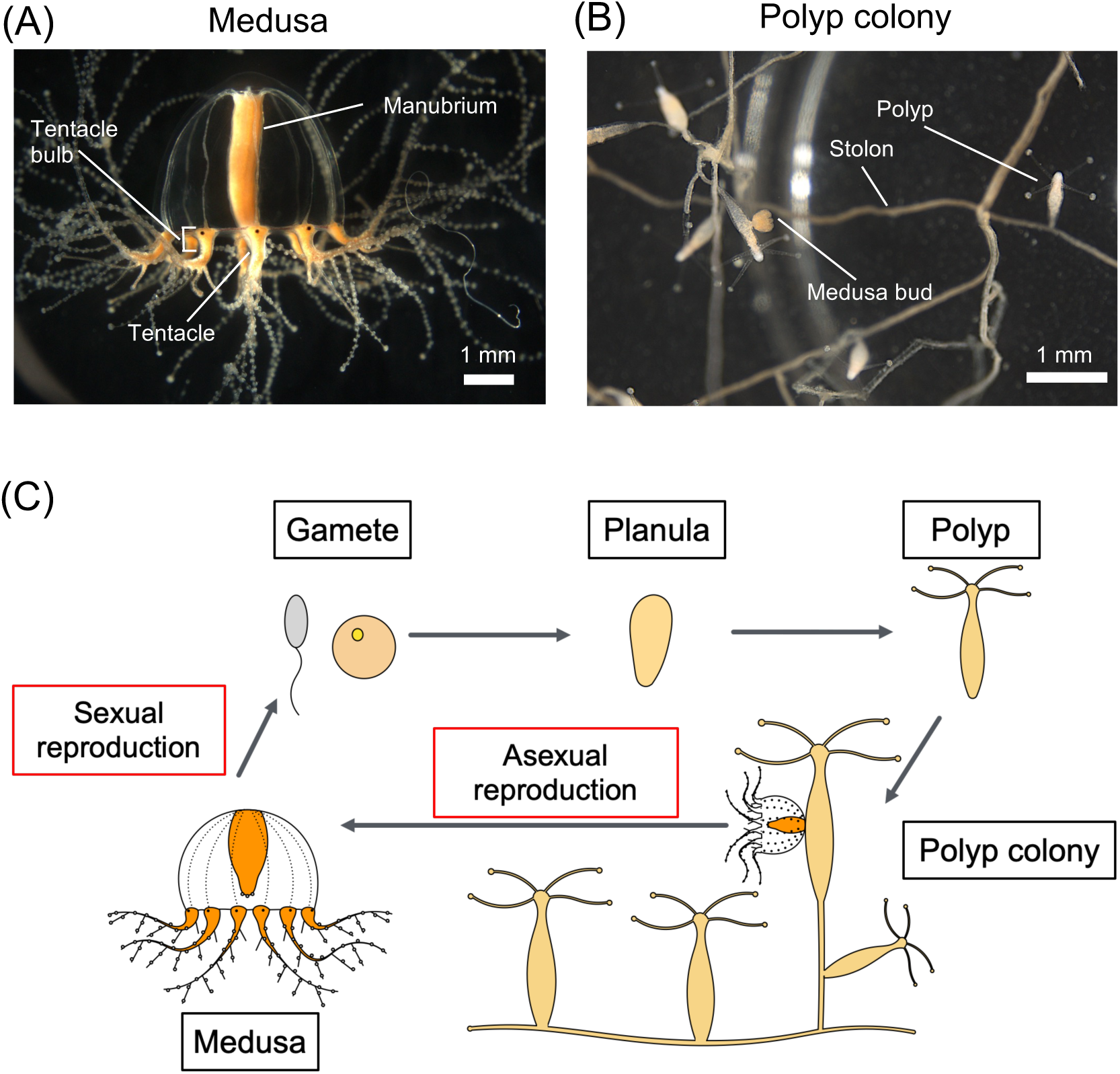
Appearance and life cycle of *C. pacificum*. (A) Lateral view of an adult medusa (strain 6W). Branching tentacles are a characteristic of *C. pacificum* (B) Polyp colony where polyps are interconnected by stolons. (C) Schematic illustration of the life cycle of *C. pacificum*.

In this study, we generated the first chromosome-level genome assembly for *C. pacificum*. K-mer analysis indicates high heterozygosity in the sequenced 6W strain, with an estimated rate of 3.89%. The primary and secondary assemblies span approximately 504 Mb and 552 Mb, respectively. This genomic resource expands the repertoire of high-quality hydrozoan genomes and provides a foundation for comparative genomic analyses of genome organization, synteny conservation, and genome evolution across Cnidaria. It will also facilitate future molecular and functional studies using *C. pacificum* as an emerging experimental model for regeneration, development, physiology, and behavior.

## Methods

### Sample collection

The hydrozoan jellyfish *C. pacificum* (Fig. 1) has been maintained as a laboratory model since its establishment by Deguchi and colleagues^13^. Two laboratory strains, routinely maintained as asexually propagating polyp colonies, were used in this study. Because each strain is maintained through clonal propagation of polyp colonies, individuals within each strain are expected to share a largely homogeneous genetic background. The female strain 6W was established in 2002 from wild individuals collected at Orinohama Fishing Port, Ishinomaki City, Miyagi Prefecture, Japan, whereas the male strain UN2 was established in 2012 from wild individuals collected at Nonoshima Island, Shiogama City, Miyagi Prefecture, Japan^28^. Both strains were transferred to our laboratory in 2017 and have been maintained under standard laboratory conditions since then.

Medusae were maintained at 22 °C under a 23.5 h light/0.5 h dark photoperiod, whereas polyp colonies were maintained at 22 °C under constant darkness. Animals were cultured in 24-ppt artificial seawater (ASW; SEA LIFE, Marin Tech) prepared with dechlorinated tap water and maintained in plastic containers (V-7, V-8, and V-9; AS ONE). Medusae and polyps were fed newly hatched *Artemia* nauplii (A&A Marine LLC).

Genome sequencing was performed using medusae from the female strain 6W. Some RNA-seq datasets used for gene prediction were generated from the male strain UN2 (Table S1).

### Library construction and sequencing

Sequencing libraries for PacBio HiFi genome sequencing and Hi-C sequencing were prepared from medusae of the female strain 6W (Fig. 2A). High-molecular-weight genomic DNA (HMW gDNA) was extracted from 40 one-month-old frozen medusae using a Genomic-tip 20/G kit (QIAGEN). The 40 medusae were asexually propagated and considered genetically clonal. Briefly, two batches of 20 medusae were pulverized under liquid nitrogen, lysed in G2 buffer supplemented with 2-mercaptoethanol, proteinase K and RNase A, and purified using Genomic-tip columns followed by AMPure XP (Beckman Coulter) bead cleanup. DNA quality and concentration were assessed using a Qubit 2.0 Fluorometer (Thermo Fisher Scientific), a NanoDrop 2000 spectrophotometer (Thermo Fisher Scientific), and pulsed-field gel electrophoresis (CHEF Mapper, Bio-Rad).

**Figure 2.**
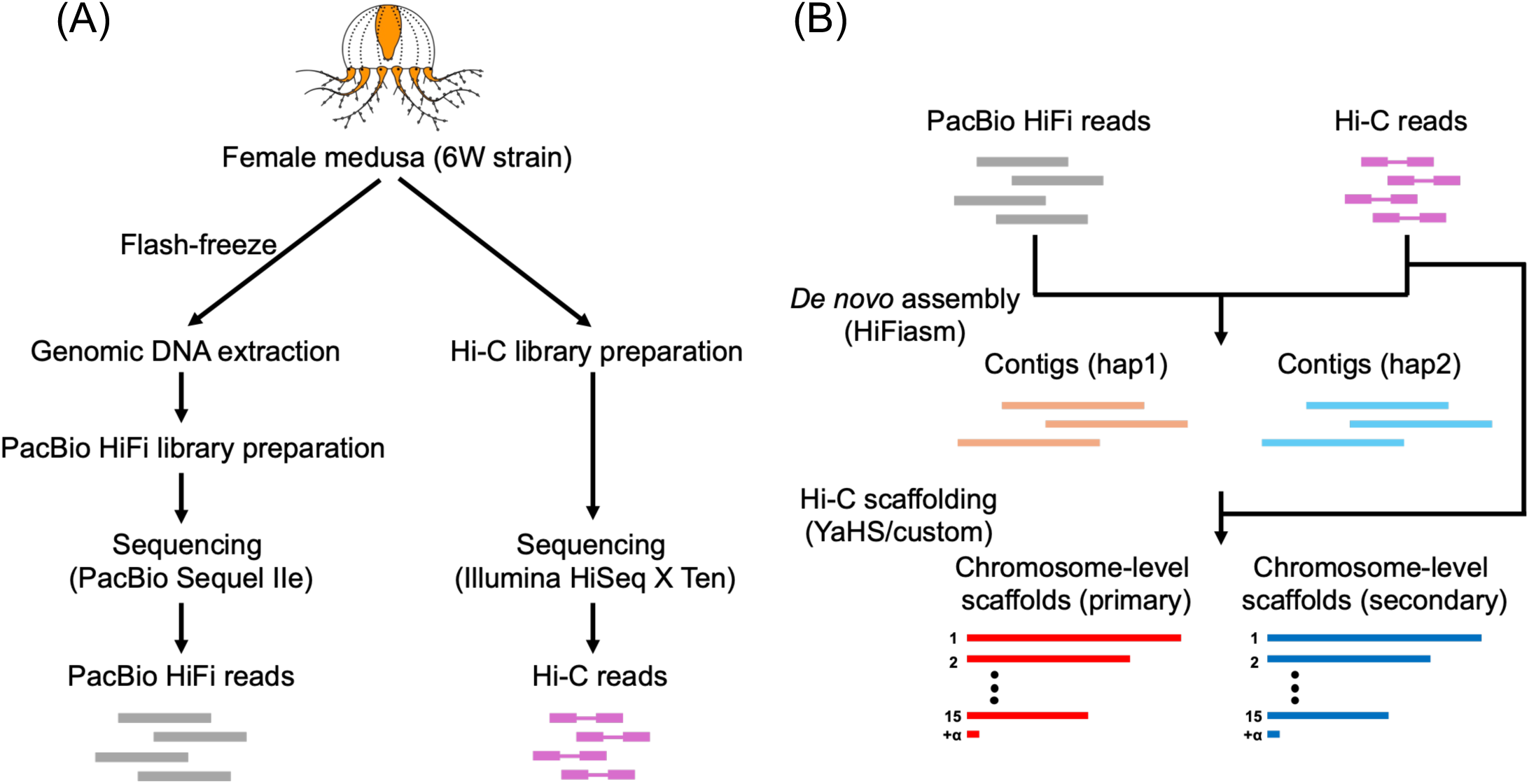
Experimental design and genome assembly pipeline. (A) Overview of the experimental workflow and genome assembly pipeline of *C. pacificum* in this study. (B) Flowchart illustrating the chromosome-level genome assembly of *C. pacificum*.

For PacBio HiFi sequencing, 5.0 μg of HMW gDNA was sheared to 15–20 kb using a Megaruptor 3 system (Diagenode). A SMRTbell library was constructed using the SMRTbell Express Template Prep Kit 2.0 (Pacific Biosciences) according to the manufacturer’s instructions and size-selected using the BluePippin system (Sage Science) with a 0.75% agarose cassette and a high-pass 6–10 kb protocol. The final library had an estimated insert size of 22 kb and a concentration of 33.3 ng/µL. Sequencing was performed on a PacBio Sequel IIe platform at the National Institute for Basic Biology using Circular Consensus Sequencing (CCS) mode with a 30-h movie time, and HiFi reads were generated using SMRT Link v11.0. A total of 1,661,872 HiFi reads were obtained, corresponding to 30.33 Gb of sequence data, with a read N50 of 18.5 kb (Table 1).

**Table 1.** Sequencing datasets generated and used in this study.

For chromosome conformation capture, an Omni-C proximity ligation library was prepared from 19 live medusae using the Omni-C Proximity Ligation Assay kit (Dovetail Genomics) according to the manufacturer’s protocol for non-mammalian samples. Briefly, medusae were washed with seawater, and the amount of extraction buffer was increased to reduce the potential effect of seawater carryover. After chromatin crosslinking, the sample was subjected to nuclease digestion, proximity ligation, DNA purification and Illumina library preparation. Chromatin digestion efficiency was estimated to be 70%, and 382 ng of DNA was recovered after proximity ligation. A total of 150 ng of DNA was used for library preparation, followed by ligation capture and 12 cycles of PCR amplification. The insert size distribution of the resulting library was evaluated using an Agilent 2100 Bioanalyzer (Agilent Technologies), and the final Omni-C library had an average fragment size of 935 bp and a total yield of 1,086 ng. The Omni-C library was sequenced by Macrogen Japan using an Illumina HiSeq X platform with paired-end 150-bp reads. In total, 821 million reads, corresponding to 124.0 Gb of raw sequence data, were obtained (Table 1).

To obtain transcriptomic evidence for gene annotation, we compiled six previously generated RNA-seq datasets representing diverse tissues, developmental stages, and experimental conditions (Table 1; Table S1). The first dataset comprised ectoderm-derived epithelial tissue, oocytes, and manubrium from medusae of the 6W strain (unpublished dataset provided by Dr. Takeda). The second dataset was a previously published RNA-seq dataset^26^ generated from ring tissue and manubrium of medusae from the UN2 strain. The third dataset consisted of samples collected during tentacle regeneration and following DAPT or xyloside treatment in the 6W strain. The fourth dataset comprised polyp and stolon samples collected during dormancy and recovery from dormancy in the UN2 strain. The fifth dataset consisted of dormant and recovering stolon samples treated with a TRPM inhibitor or DMSO control. The sixth dataset comprised medusa buds and medusa bud-derived cysts from the 6W strain.

### Iso-Seq transcriptomic sequencing across developmental stages and body forms

To complement the short-read RNA-seq datasets and provide long-read transcriptomic evidence for gene prediction, we generated two PacBio Iso-Seq datasets spanning multiple developmental stages and body forms of *C. pacificum*. For the first dataset, total RNA was extracted from embryos, planulae, primary polyps, medusa buds, cysts derived from medusa buds and polyps, polyps, stolons, and medusae using the RNeasy Micro Kit (QIAGEN), followed by purification with the RNA Clean & Concentrator-5 kit (Zymo Research). Embryos, planulae, and primary polyps were derived from crosses between the 6W and UN2 strains, whereas the medusa material comprised individuals from both strains. RNA obtained from the different developmental stages and body forms was pooled and submitted to Novogene for PacBio Iso-Seq library preparation and sequencing. At Novogene, cDNA synthesis was performed using the Iso-Seq Express 2.0 Kit, followed by library preparation using the Kinnex Full-Length RNA Kit. The resulting full-length non-concatemer (FLNC) read dataset comprised 16,928,185 reads totaling 14.12 Gb, with a mean read length of 834.2 bp (Table 1).

For the second dataset, total RNA was extracted separately from six samples representing embryos, planulae, stolons, polyps, and two medusa samples, using TRIzol reagent (Thermo Fisher Scientific), followed by purification with the RNA Clean & Concentrator-5 kit (Zymo Research). The embryo and planula samples were derived from crosses between the 6W and UN2 strains, whereas the medusa samples included individuals from both strains. For each sample, 300 ng of total RNA was used for cDNA synthesis using the Iso-Seq Express 2.0 Kit (Pacific Biosciences). The six samples were individually barcoded using Iso-Seq primers bc02–bc07 and processed using the Kinnex Full-Length RNA workflow. The resulting library was sequenced on a PacBio Sequel IIe system. The resulting FLNC read dataset comprised 5,421,863 reads totaling 12.11 Gb, with a mean read length of 2,233.1 bp (Table 1). Together, these datasets provide long-read transcript coverage across diverse developmental stages and body forms of *C. pacificum*, complementing the short-read RNA-seq datasets used for gene annotation.

### *De novo* genome assembly and Hi-C scaffolding

Genome size and heterozygosity were estimated from the PacBio HiFi reads of the *C. pacificum* 6W strain using k-mer analysis. Canonical 25-mers were counted using Jellyfish v2.3.1^29^ with the -C option. A k-mer frequency histogram was generated using jellyfish histo with the -h 100000000 option and analyzed using GenomeScope2 v2.0.1^30^ with a diploid model (p = 2), k = 25, and otherwise default parameters. The estimated haploid genome size was 519.4 Mb, and the estimated heterozygosity was 3.89% (Supplementary Fig. S1), indicating high heterozygosity in the sequenced 6W strain. PacBio HiFi long reads were assembled using hifiasm v0.19.5^31–33^, with the Hi-C integration mode enabled (Fig. 2B). Two haplotype-resolved contig assemblies (hap1 and hap2) were generated and concatenated into a single FASTA file for chromosome-scale scaffolding with YaHS v1.2a.1^34^ (Table 2). The GC content of both assemblies was approximately 32%, indicating that the *C. pacificum* genome is AT-rich, as reported for other hydrozoans^8,35^. Throughout this study, each FASTA sequence entry was regarded as a scaffold, whereas contigs were defined as contiguous sequences separated by stretches of Ns.

**Table 2.** Statistics of the *C. pacificum* genome assemblies.

The merged assembly was indexed using SAMtools v1.19.2^36^, and a genome-size file was generated from the FASTA index. The assembly was subsequently indexed using BWA v0.7.17-r1188^37^. Paired-end Hi-C reads were aligned to the merged haplotype contig assembly using bwa mem with the −5SP and -T0 options. The resulting alignments were processed using pairtools v1.0.3^38^. Briefly, alignments were parsed using a minimum mapping-quality threshold of 40, sorted, deduplicated, and split into pairs and SAM outputs. The SAM output was converted to BAM format and sorted using SAMtools. The resulting deduplicated and sorted BAM file was indexed and used as the Hi-C alignment input for YaHS. YaHS was then run using the merged haplotype contig assembly and the processed Hi-C BAM file to generate chromosome-scale scaffolds.

To evaluate the scaffolding results, the juicer pre utility implemented in YaHS v1.1 was applied to the YaHS binary output, the final AGP file, and the FASTA index of the merged assembly. Juicer Tools v2.20.00^39^ was then used to generate a Hi-C contact map from the sorted alignments and the scaffold-size file. The contact map was visualized using Juicebox^40^ (Fig. 3) and manually inspected to infer putative allelic scaffold pairs.

**Figure 3.**
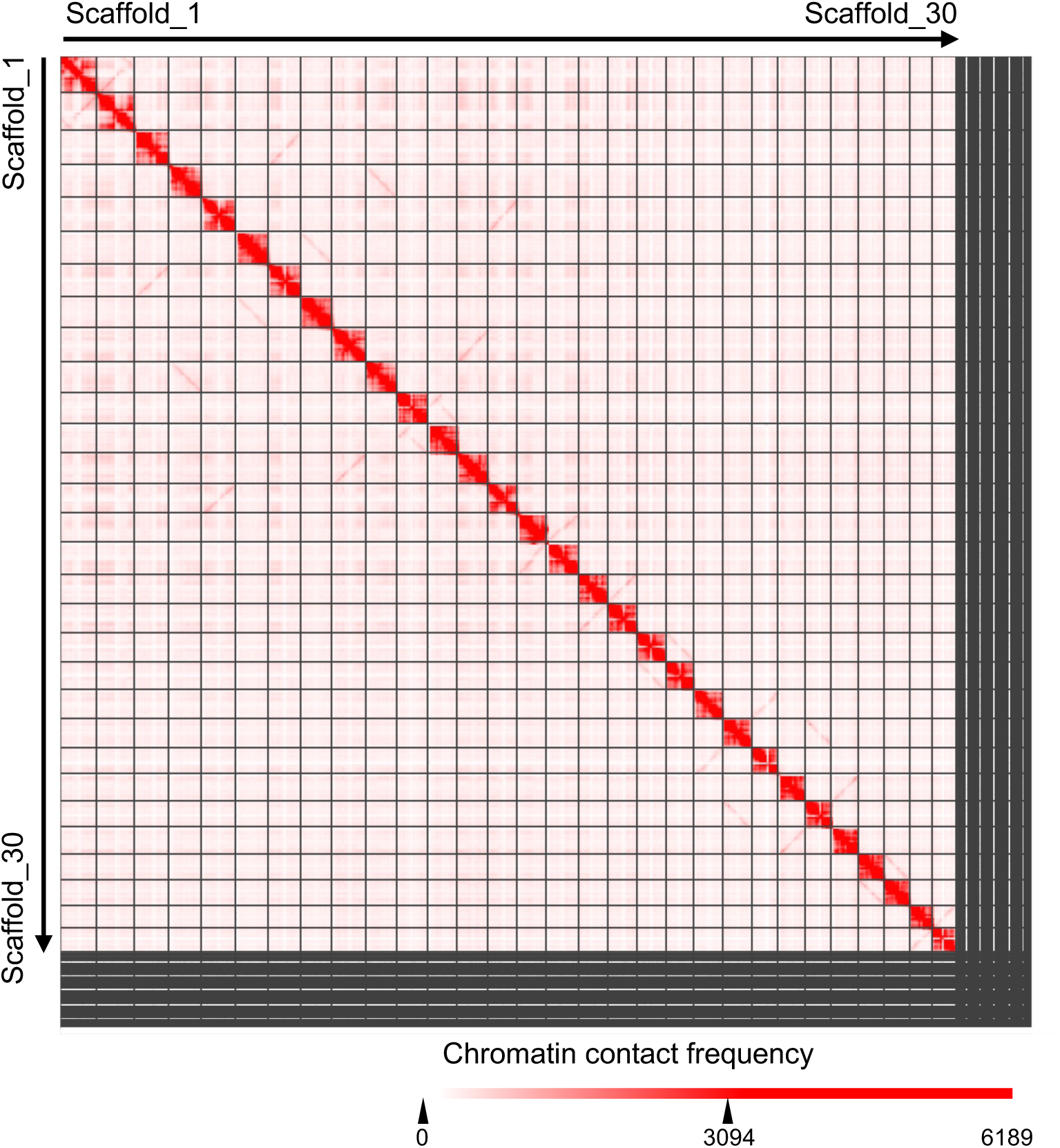
Hi-C contact map of the scaffolded *C. pacificum* assembly. The map shows the combined scaffold set before selection of the primary and secondary assemblies. Color intensity indicates chromatin contact frequency. Thirty chromosome-scale scaffolds were assigned to 15 homologous pairs based on sequence alignments and manual inspection of the contact map.

To identify homologous chromosome pairs, all-versus-all sequence alignments of the assembled scaffolds were performed using MUMmer v4.0.1^41^. Based on the sequence-alignment results and manual inspection of the Hi-C contact map, the 30 large scaffolds were assigned to 15 homologous pairs, which likely represent the 15 major chromosomes of the haploid genome. This chromosome-level organization is consistent with an inferred diploid chromosome number of 2n = 30. A haploid chromosome number of n = 15 has also been reported in diverse cnidarian species, including *Hydra*, *Nematostella*, *Hydractinia*, and *Clytia*^3,8,9,42,43^. The recovery of 15 chromosome-scale scaffold pairs in *C. pacificum* is therefore consistent with the broadly conserved chromosome number observed across these cnidarian lineages and provides additional support for the chromosome-scale scaffolding.

To construct a representative haploid sequence set, one scaffold was selected from each of the 15 homologous chromosome pairs. Scaffold selection was based on two criteria. First, the scaffold containing the greater number of complete BUSCO genes was selected. The completeness of the genome assemblies was evaluated using BUSCO v5.8.2^44^. When both scaffolds contained the same number of BUSCO genes, the longer scaffold was selected. The shorter scaffolds were subsequently evaluated for redundancy based on the output of purge_dups, and 121 nonredundant shorter scaffolds were incorporated into the representative haploid sequence set.

The resulting primary assembly consisted of 15 major chromosome-level scaffolds and 121 shorter scaffolds, with a total length of 503,996,576 bp and a scaffold N50 of 32.1 Mb (Table 2). The cumulative length of the 15 chromosome-level scaffolds was approximately 482 Mb, indicating that most of the primary assembly was represented by chromosome-level sequences. For supplementary and comparative analyses, the alternative scaffold from each of the 15 homologous chromosome pairs was combined with the remaining sequences to construct a secondary assembly. The secondary assembly comprised 551 sequences, with a total length of 551,650,960 bp and a scaffold N50 of 32.3 Mb. The 15 chromosome-level scaffolds in the secondary assembly had a cumulative length of approximately 489 Mb. Thus, most of both the primary and secondary assemblies was represented by chromosome-level scaffolds. Relative to the k-mer-based haploid genome size estimate of 519.4 Mb, the primary and secondary assemblies were approximately 3.0% smaller and 6.2% larger, respectively.

The completeness of the primary and secondary genome assemblies was evaluated using BUSCO with the Cnidaria-specific cnidaria_odb12 lineage dataset and genome mode (-m genome). In the primary assembly, 2,682 of the 3,203 BUSCO orthologs (83.7%) were identified as complete. These included 2,627 complete single-copy BUSCOs (82.0%) and 55 complete duplicated BUSCOs (1.7%). A further 234 BUSCOs (7.3%) were fragmented, whereas 287 BUSCOs (9.0%) were missing (Table 3). In the secondary assembly, 2,692 BUSCOs (84.0%) were complete, comprising 2,626 complete single-copy BUSCOs (82.0%) and 66 complete duplicated BUSCOs (2.1%). In addition, 225 BUSCOs (7.0%) were fragmented and 286 BUSCOs (8.9%) were missing (Table 3). The representative haploid assembly was subsequently used as the reference genome for gene prediction, functional annotation, transcriptomic analyses, and comparative genomic analyses.

**Table 3.** BUSCO assessment of the genome assembly and gene annotation.

### Analysis of repetitive elements

To identify repetitive elements in the *C. pacificum* genome, a custom repeat library was generated using RepeatModeler v2.0.6^45^ with the default parameters (Fig. 4A, 4B). RepeatMasker v4.1.7-p1^46^ was used to detect repetitive elements in the genome assembly to produce a soft-masked genome sequence. The repeat landscape was generated using calcDivergenceFromAlign.pl and createRepeatLandscape.pl provided with RepeatMasker, and the genomic proportions of individual repeat classes were visualized based on the RepeatMasker summary table (.tbl). A total of 265 Mb (52.52%) of the genome sequences were masked by RepeatMasker as repetitive elements and low complexity regions (Fig. 4B). Among the classified transposable elements, LINEs (Long INterspersed Elements) (4.44%), LTRs (Long Terminal Repeats) (2.00%), and DNA transposons (4.14%) were the major components (Fig. 4B; Table 4). The proportion of repetitive elements in the *C. pacificum* genome (52.52%) is similar to other hydrozoans, such as *Hydra vulgaris* AEP (65.3%)^6^ and *Hydractinia symbiolongicarpus* (61%)^8^.

**Figure 4.**
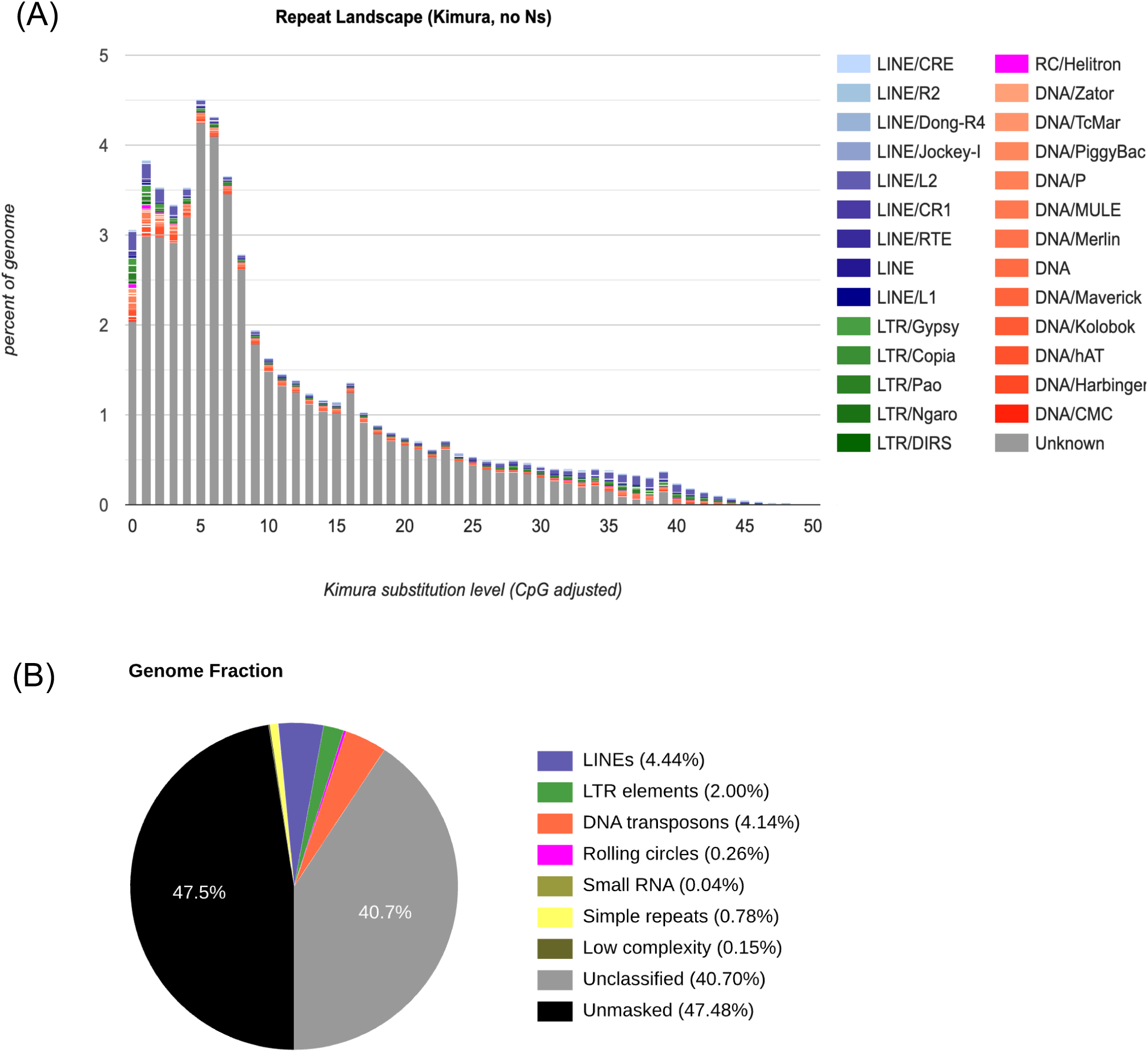
Repetitive elements in the *C. pacificum* genome. (A) Repeat landscape of the *C. pacificum* genome. The x-axis indicates the Kimura substitution level (%), representing sequence divergence from repeat consensus sequences, and the y-axis shows the proportion of the genome occupied by each repeat class. (B) Genome composition based on the RepeatMasker summary table. Percentages were calculated relative to the genome length excluding N/X-runs.

**Table 4.** Summary of repetitive elements identified in the *C. pacificum* genome.

### Gene prediction and functional annotation

To predict genes in the *C. pacificum* genome assembly, we used BRAKER3^47–59^, integrating ab initio gene prediction with RNA-seq alignments, Iso-Seq-derived hints, and protein homology evidence. RNA-seq reads from the six transcriptome datasets (Table S1) were aligned to the reference genome using STAR v2.7.10b^60^ with default parameters, and the resulting BAM files were merged using SAMtools. To complement the transcriptomic evidence, two PacBio Iso-Seq datasets were processed using the Iso-Seq pipeline^61^ provided by Pacific Biosciences. Full-length non concatemer (FLNC) reads were generated with lima v2.12.0 and isoseq v4.2.0, and subsequently clustered using isoseq cluster2. The resulting clustered isoforms were aligned to the genome using pbmm2 v1.16.99 and collapsed using isoseq collapse. Exons extracted from the collapsed GFF files were supplied to BRAKER3 using the “--hints” option. Protein homology evidence was obtained from the metazoan protein dataset in OrthoDB v12^44^ (Table S1). Gene prediction was performed on the soft-masked genome using the “--softmasking --gff3” options, resulting in the prediction of 17,206 protein-coding genes.

Completeness of the predicted gene set was evaluated using BUSCO with the cnidaria_odb12 lineage dataset in protein mode. To generate a non-redundant protein set containing one representative protein per gene, the longest protein isoform for each predicted gene was extracted using SeqKit v2.3.0^62^ and seqtk v1.4-r122^63^. A total of 2,833 (88.4%) complete BUSCO genes were identified, including 2,767 (86.4%) single-copy and 66 (2.1%) duplicated BUSCOs, whereas 100 (3.1%) BUSCOs were fragmented and 270 (8.4%) were missing (Table 3).

Functional annotation of the predicted 17,206 proteins was performed using BLASTP v2.16.0^64^ with the parameter “-evalue 1e-20” against Swiss-Prot^65^. In total, 8,677 (50.43%) predicted proteins had significant hits (Table 5). Protein domains were annotated using InterProScan v5.76-107.0^66^, resulting in annotations for 16,346 (95.00%) predicted proteins (Table 5). Orthology-based gene functional annotation was performed using eggNOG-mapper v2.1.13^67^, which annotated 12,803 (74.41%) predicted proteins (Table 5). Overall, 16,470 (95.72%) of the predicted proteins were assigned at least one functional annotation (Fig. 5A; Table 5).

**Figure 5.**
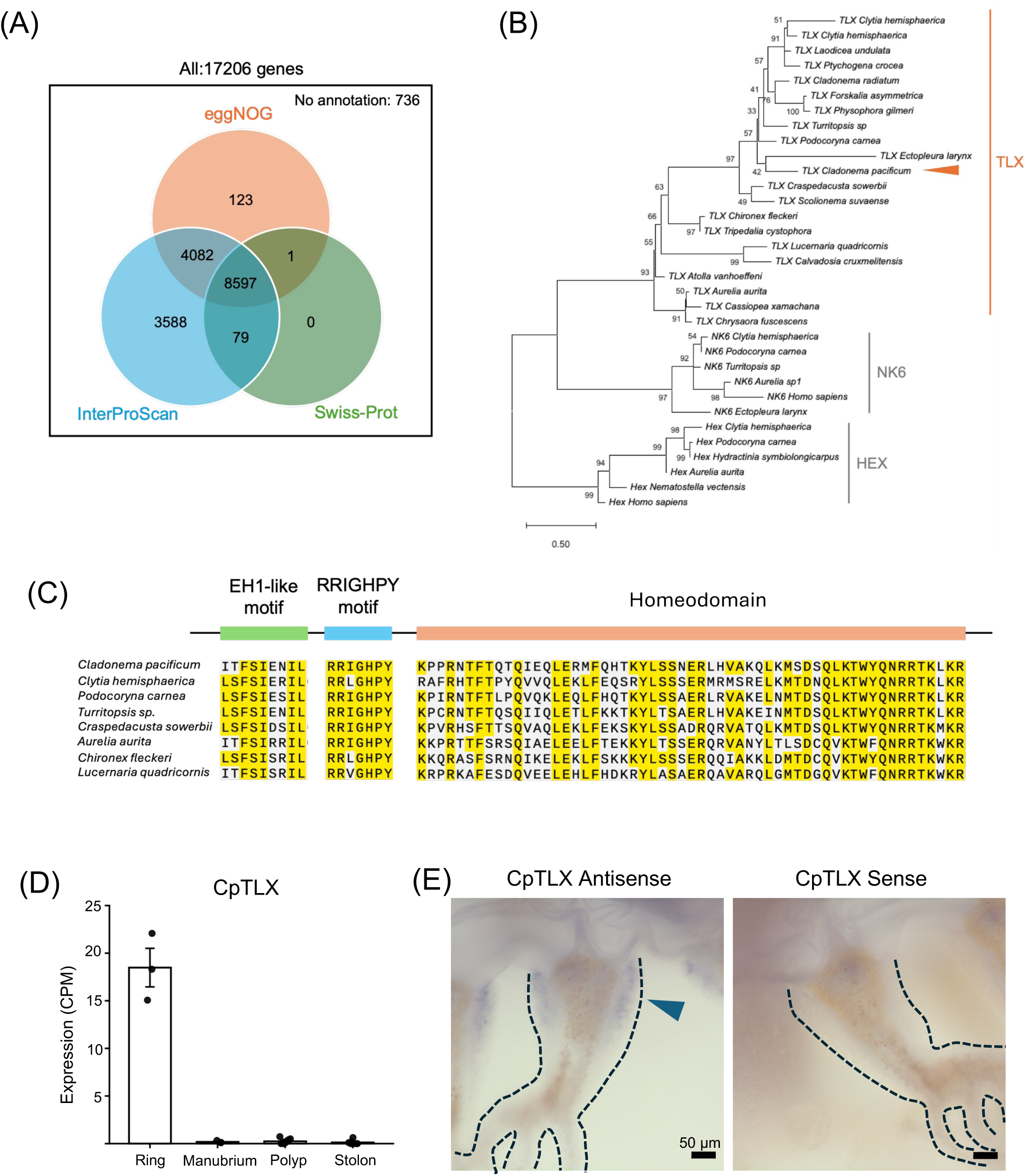
Functional annotation and locus-specific validation using TLX. (A) Venn diagram summarizing functional annotation of predicted protein-coding genes using Swiss-Prot, InterProScan, and eggNOG-mapper. (B) Maximum-likelihood phylogeny of TLX, NK6, and Hex homeobox proteins. The *C. pacificum* TLX candidate (Cpac_g3151) clusters within the cnidarian TLX clade. (C) Conserved domain organization and sequence alignment of the *C. pacificum* TLX protein compared with representative cnidarian TLX proteins. (D) Expression of Cpac_g3151 quantified from RNA-seq datasets. Ring samples correspond to the umbrella rim including the tentacle bulbs. Data are presented as mean ± SEM. Each dot represents an individual biological replicate. (E) Whole-mount chromogenic *in situ* hybridization of Cpac_g3151. The antisense probe detected expression in the tentacle bulb (arrowhead), whereas the sense probe served as a negative control. n=4 each.

**Table 5.** Functional annotation of predicted protein-coding genes.

### Locus-specific validation of genome annotation using TLX

To provide locus-specific technical validation of the genome annotation and associated transcriptomic datasets, we selected the TLX homeobox gene as a representative case study. TLX has previously been implicated in medusa development in cnidarians^68^. A candidate TLX ortholog (Cpac_g3151) was identified from the BRAKER3 gene models as the top hit and was independently annotated as a homeobox-containing transcription factor by both eggNOG-mapper and InterProScan.

To evaluate the orthology of this candidate, amino acid sequences of cnidarian TLX proteins together with representative NK6 and Hex proteins (Table S2) were aligned using MAFFT v7.505^69^. The resulting alignment was trimmed using trimAl v1.5.1^70^, and maximum-likelihood phylogenetic inference was performed using IQ-TREE v3.0.1^71^. The *C. pacificum* sequence clustered within the cnidarian TLX group and was clearly separated from NK6 and Hex, supporting its assignment as a TLX ortholog candidate (best fit model: Q.INSECT+G4) (Fig. 5B). Because the tree was inferred from conserved regions, internal branching relationships within the TLX clade were not interpreted further. The predicted TLX gene model was manually inspected using IGV v2.16.0^72^. RNA-seq read coverage upstream of the annotated first exon suggested that the original gene model was incomplete. An upstream genomic region was therefore examined for alternative translation initiation sites, and the coding sequence was manually extended based on continuous RNA-seq coverage and open reading frame prediction. This refined sequence recovered an N-terminal EH1-like motif candidate (ITFSIENIL) in addition to the conserved RRIGHPY motif and homeodomain. The extended *C. pacificum* TLX sequence was comparable to other cnidarian TLX proteins in retaining the EH1-like motif candidate, the RRIGHPY motif, and the homeodomain (Fig. 5C).

To examine expression of the refined TLX gene model, RNA-seq reads from medusa and non-medusa tissues were quantified using featureCounts implemented in Subread v2.1.1^73^. Gene expression levels were calculated as counts per million (CPM) and compared among tissues. The TLX candidate showed markedly elevated expression in jellyfish ring samples (mean CPM = 18.48), whereas expression was very low in jellyfish manubrium (mean CPM = 0.18), polyp (mean CPM = 0.24), and stolon (mean CPM = 0.12) samples (Fig. 5D).

### Experimental validation of TLX expression by whole-mount *in situ* hybridization

To examine the spatial expression pattern of the candidate TLX ortholog (Cpac_g3151), DIG-labeled RNA probes were generated from a partial coding sequence amplified from cDNA prepared from medusae of the UN2 strain using Ex Taq HS (Takara Bio). The PCR product was cloned into the pTA2 vector using the Target Clone TA Cloning Kit (TOYOBO), and the insert sequence was confirmed by Sanger sequencing. The following primers were used for amplification:

Forward: 5′-AATGGAAGATCGAACCTCAAACT-3′
Reverse: 5′-ACCTCGATCGTTGTTCATTGAG-3′

Sense and antisense RNA probes were synthesized by *in vitro* transcription using T7 or T3 RNA polymerase and DIG RNA Labeling Mix (Roche). Linear DNA templates were generated by PCR using M13 universal primers (M13-21: 5′-TGTAAAACGACGGCCAGT-3′; M13-Rev: 5′-GAGCGGATAACAATTTCACACAGG-3′). DIG-labeled RNA probes were purified using ProbeQuant G-50 Micro Columns (Cytiva), mixed with an equal volume of formamide, and stored at −30 °C until use. Sense probes were used as negative controls.

Adult medusae of the UN2 strain were anesthetized for 10 min in ASW containing 7% MgCl₂ and fixed in 4% paraformaldehyde for 1 h at room temperature or overnight at 4 °C. Whole-mount chromogenic *in situ* hybridization was performed essentially as described previously for *C. pacificum*^16^, with minor modifications. Following fixation, samples were washed in PBST (PBS containing 0.1% Tween-20), pre-hybridized for 2 h at 55 °C, and hybridized overnight (21 h) at 55 °C with DIG-labeled RNA probes (0.5–1.0 ng μL⁻¹). After a series of post-hybridization washes with SSC/formamide buffers, samples were blocked in 1% blocking reagent (Roche) and incubated overnight at 4 °C with alkaline phosphatase-conjugated anti-DIG antibody (Roche; 1:2,000). Color development was performed using NBT/BCIP substrate solution for approximately 8 h in the dark and terminated by washing with TNT buffer. Samples were mounted in 70% glycerol and imaged using a Zeiss Axio Zoom microscope.

Whole-mount chromogenic *in situ* hybridization revealed strong expression of TLX in the tentacle bulb of adult medusae (Fig. 5E), consistent with the medusa-associated expression pattern identified by RNA-seq analysis (Fig. 5D). Together, these analyses support the assignment of Cpac_g3151 as a TLX ortholog and illustrate the utility of the genome assembly and associated transcriptomic datasets for ortholog identification, gene model refinement, and gene expression analysis.

### Molecular phylogenetic analysis of *C. pacificum* within Cnidaria

*C. pacificum* is taxonomically classified within the phylum Cnidaria, class Hydrozoa, order Anthoathecata, and family Cladonematidae^14,74^. To examine its phylogenetic position using genome-scale data, we performed a phylogenomic analysis, including *C. pacificum* and ten additional representative cnidarian species (Table S3). Genome assemblies and gene annotations were obtained from the NCBI Genome database. Protein sequences were extracted using GffRead v0.12.7^55^, and the longest protein isoform for each gene was retained.

Orthologous gene families were identified using OrthoFinder v3.1.4^75^, yielding 19,459 orthogroups. A total of 2,847 single-copy orthogroups were aligned using MAFFT, and poorly aligned regions were removed using GBLOCKS v1.0^76,77^ with the parameters “-t=p -b1=6 -b2=6 -b3=8 -b4=10 -b5=n”. Alignments shorter than 50 amino acids after trimming were excluded. The remaining alignments were concatenated using AMAS.py v1.0^78^ to generate a supermatrix.

Maximum-likelihood (ML) phylogenetic inference was performed using IQ-TREE with ModelFinder and branch support estimated using 1,000 ultrafast bootstrap replicates and 1,000 SH-aLRT replicates (-m MFP -B 1000 --alrt 1000). The resulting ML tree was visualized using MEGA12^79^ and placed *C. pacificum* within Hydrozoa with strong bootstrap support (Fig. 6A).

**Figure 6.**
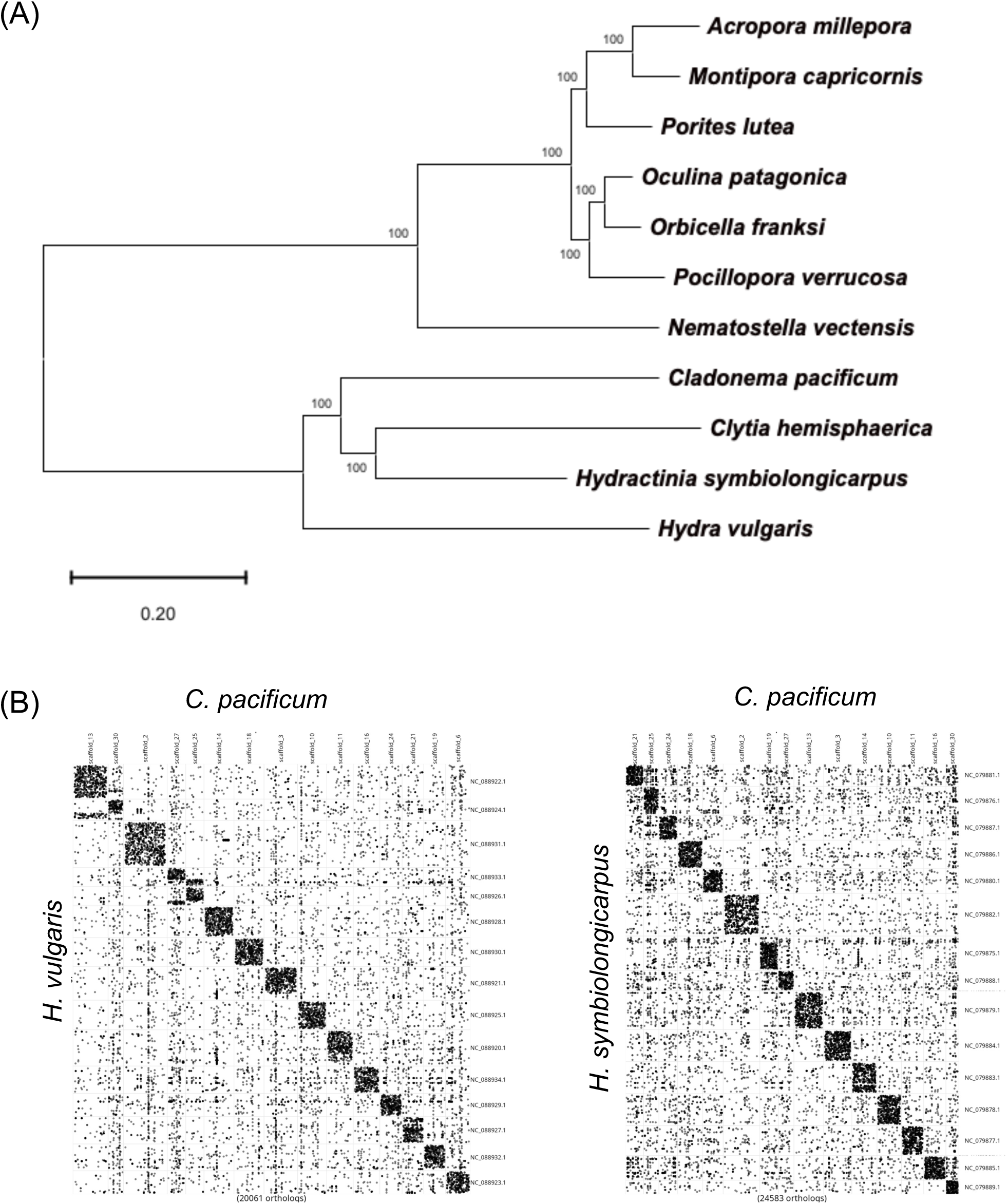
Comparative genomic analyses of *C. pacificum* and chromosomal synteny analyses. (A) Maximum-likelihood phylogeny of *C. pacificum* and ten representative cnidarian species inferred from 2,847 single-copy orthogroups. Branch support values were estimated from 1,000 bootstrap replicates. Scale bar indicates amino acid substitutions per site. (B) Oxford plots showing chromosome-scale synteny between *C. pacificum* and *Hydra vulgaris* (left), and between *C. pacificum* and *Hydractinia symbiolongicarpus* (right). Each dot represents an orthologous gene pair.

### Chromosomal synteny analysis

Pairwise macrosyntenic relationships between *C. pacificum* and two representative hydrozoans, *Hydra vulgaris* and *Hydractinia symbiolongicarpus*, were examined using the R package macrosyntR^80^. Pairwise ortholog assignments inferred by OrthoFinder were used as input, and Oxford plots were generated to visualize conservation of chromosomal linkage between genomes (Fig. 6B). The resulting plots revealed broadly conserved chromosome-scale synteny between *C. pacificum* and both hydrozoan species.

## Data Records

All raw sequencing data produced in this study have been deposited in the DDBJ Sequence Read Archive (DRA). The PacBio HiFi, Hi-C, and PacBio Iso-Seq sequencing data were deposited under BioProject accession PRJDB42964^81^. The PacBio HiFi and Hi-C reads are available under accession numbers DRR1271270 and DRR1271269, respectively.

The RNA-seq datasets were deposited under BioProject accessions PRJDB45818 (ectodermal epithelium, oocytes, and manubrium)^82^, PRJDB42990 (stolon samples treated with a TRPM inhibitor)^83^, PRJDB42987 (polyp and stolon dormancy and recovery)^84^, PRJDB42972 (tentacle regeneration)^85^, and PRJDB42991 (medusa buds and medusa bud-derived cysts)^86^. All RNA-seq and PacBio Iso-Seq datasets used for BRAKER3 gene prediction, including previously published datasets, are listed in Supplementary Table S1 together with their accession numbers.

The primary genome assembly has been deposited in the DNA Data Bank of Japan (DDBJ) under WGS accession numbers BAANGU010000001-BAANGU010000136^87^.

The primary and secondary genome assemblies, a scaffold-to-accession mapping table for the primary assembly, repetitive element annotations, the custom repeat library, gene models, protein and coding sequences, functional annotations, and other related materials have been deposited in Figshare^88^. The gene models and repetitive element annotations are based on the primary genome assembly. Functional annotations were generated using the longest protein isoform of each gene. The manually extended TLX sequence is provided separately.

## Technical Validation

The quality of the gDNA and RNA was evaluated in terms of purity, concentration, and size distribution to ensure high quality before sequencing. RNA-seq reads were evaluated using fastp v0.24.3^89^ prior to downstream analyses. The RNA-seq reads from the six experiments used for genome annotation were aligned to the genome using STAR, resulting in an average mapping rate of 87.33%. The completeness of the genome assembly and gene annotation was evaluated using BUSCO v5.8.2 and the Cnidaria odb12 lineage dataset (cnidaria_odb12). The BUSCO scores for the primary genome assembly and annotated protein sequences were 83.7% and 88.4%, respectively.

## Data Availability

The raw sequencing data are available in the DDBJ Sequence Read Archive under BioProject accessions PRJDB42964^81^, PRJDB45818^82^, PRJDB42990^83^, PRJDB42987^84^, PRJDB42972^85^, and PRJDB42991^86^. Accession numbers for individual sequencing datasets are provided in Supplementary Table S1 and the Data Records section. The primary genome assembly is available in DDBJ under WGS accession numbers BAANGU010000001–BAANGU010000136^87^. The primary and secondary genome assemblies, repetitive element annotations, gene models, functional annotations, and other related materials are available at Figshare^88^.

## Code Availability

All software was used according to the manuals and protocols provided by the software developers. If no detailed parameters were mentioned for the software, it was run using its default parameter settings, as described in the software documentation.

## Acknowledgements

Computations were partially performed on the NIG supercomputer at ROIS National Institute of Genetics.

## Author Contributions

J.U.: performed experiments, analyzed data and prepared figures and wrote the manuscript. K.Y.: performed experiments and analyzed data. H.N.: performed experiments and collected samples. K.N.K. and T.K.: verified the data and edited the manuscript. Y.I.: verified the data. N.T. and R.D.: collected samples and verified the data. S.S.: analyzed data, wrote, and edited the manuscript. Y.N.: supervised the whole project and experiments, prepared figures, wrote and edited the manuscript.

---

**Supplementary Figure S1.**
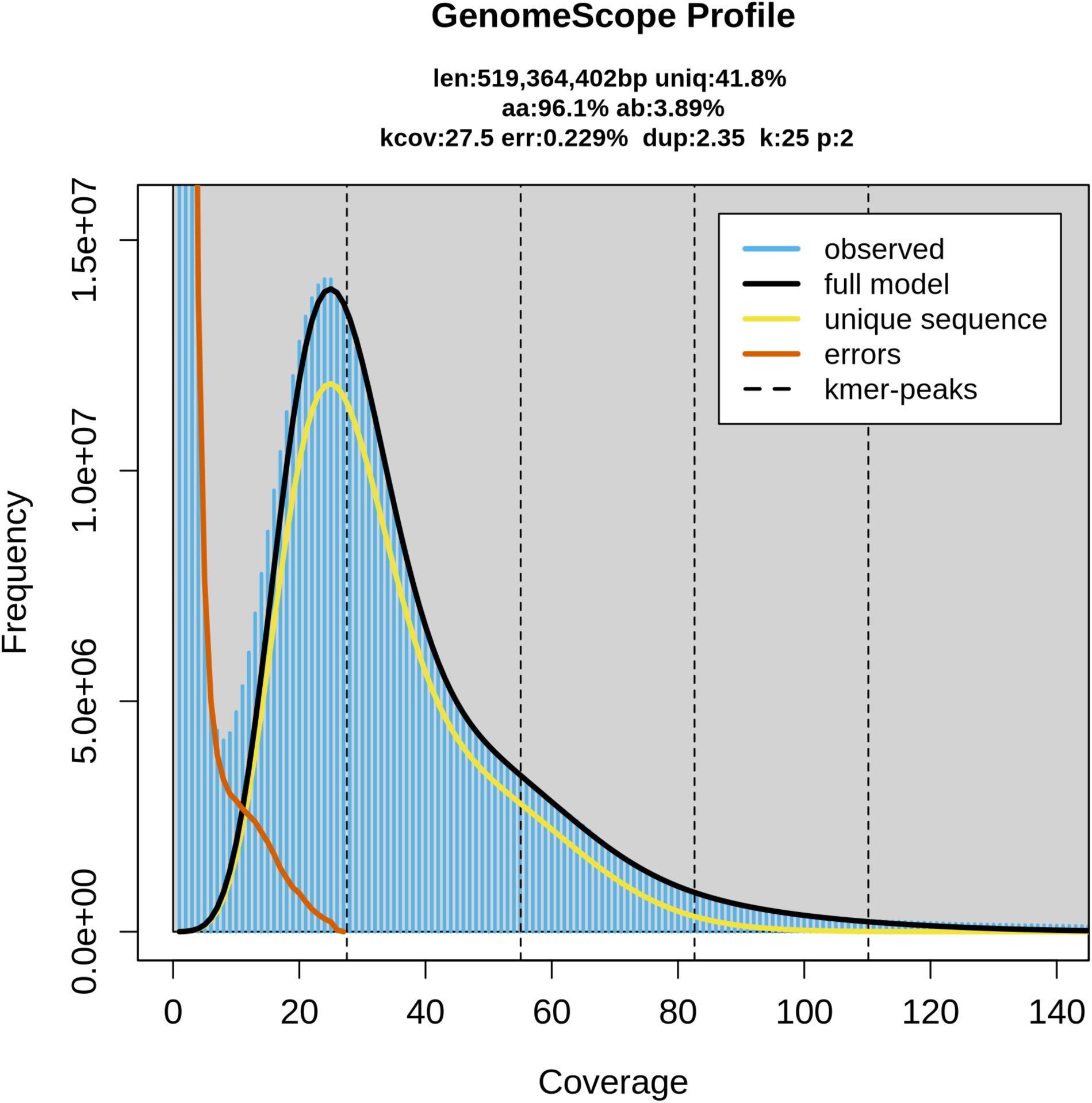
K-mer-based estimation of genome size and heterozygosity in *C. pacificum*. Canonical 25-mers were counted from PacBio HiFi reads of the 6W strain using Jellyfish v2.3.1 and analyzed using GenomeScope2 v2.0.1 with a diploid model. Observed k-mer frequencies and the fitted model are shown on linear scales. The x-axis indicates k-mer coverage, and the y-axis indicates the number of distinct k-mers. The estimated haploid genome size was 519.4 Mb, and the estimated heterozygosity was 3.89%.

## References

1. Technau, U. & Steele, R. E. Evolutionary crossroads in developmental biology: Cnidaria. Development 138, 1447–1458 (2011).

2. Zapata, F. et al. Phylogenomic Analyses Support Traditional Relationships within Cnidaria. PLOS ONE 10, e0139068 (2015).

3. Zimmermann, B. et al. Topological structures and syntenic conservation in sea anemone genomes. Nat. Commun. 14, 8270 (2023).

4. Ikmi, A., McKinney, S. A., Delventhal, K. M. & Gibson, M. C. TALEN and CRISPR/Cas9-mediated genome editing in the early-branching metazoan Nematostella vectensis. Nat. Commun. 5, 5486 (2014).

5. Vogg, M. C., Galliot, B. & Tsiairis, C. D. Model systems for regeneration: Hydra. Development 146, dev177212 (2019).

6. Kon-Nanjo, K. et al. The dynamic genomes of Hydra and the anciently active repeat complement of animal chromosomes. Genome Biol. 26, 186 (2025).

7. Frank, U., Nicotra, M. L. & Schnitzler, C. E. The colonial cnidarian Hydractinia. EvoDevo 11, 7 (2020).

8. Kon-Nanjo, K., et al. Chromosome-level genome assembly of Hydractinia symbiolongicarpus. G3 GenesGenomesGenetics 13, jkad107 (2023).

9. Leclère, L. et al. The genome of the jellyfish Clytia hemisphaerica and the evolution of the cnidarian life-cycle. Nat. Ecol. Evol. 3, 801–810 (2019).

10. Momose, T. et al. High doses of CRISPR/Cas9 ribonucleoprotein efficiently induce gene knockout with low mosaicism in the hydrozoan Clytia hemisphaerica through microhomology-mediated deletion. Sci. Rep. 8, 11734 (2018).

11. Accorsi, A., Guo, L., Marshall, W. F., Mommersteeg, M. T. M. & Nakajima, Y. Extraordinary model systems for regeneration. Development 151, dev203083 (2024).

12. Fujita, S., Kuranaga, E. & Nakajima, Y. Regeneration Potential of Jellyfish: Cellular Mechanisms and Molecular Insights. Genes 12, 758 (2021).

13. Deguchi, R., Kondoh, E. & Itoh, J. Spatiotemporal characteristics and mechanisms of intracellular Ca2+ increases at fertilization in eggs of jellyfish (Phylum Cnidaria, Class Hydrozoa). Dev. Biol. 279, 291–307 (2005).

14. Fujiki, A., Hou, S., Nakamoto, A. & Kumano, G. Branching pattern and morphogenesis of medusa tentacles in the jellyfish Cladonema pacificum (Hydrozoa, Cnidaria). Zool. Lett. 5, 12 (2019).

15. Deguchi, R. & Itoh, T. Methods for collecting and breeding the hydrozoan jellyfish Cladonema pacificum: aim for its use for academic research and education. Bull. Miyagi Univ. Educ. 40, 107–119 (2005).

16. Fujita, S., Kuranaga, E., Miura, M. & Nakajima, Y.-I. Fluorescent In Situ Hybridization and 5-Ethynyl-2’-Deoxyuridine Labeling for Stem-like Cells in the Hydrozoan Jellyfish Cladonema pacificum. J. Vis. Exp. JoVE 10.3791/64285 (2022) doi:10.3791/64285.

17. Masuda-Ozawa, T. et al. siRNA-mediated gene knockdown via electroporation in hydrozoan jellyfish embryos. Sci. Rep. 12, 16049 (2022).

18. Fujita, S., Kuranaga, E. & Nakajima, Y.-I. Cell proliferation controls body size growth, tentacle morphogenesis, and regeneration in hydrozoan jellyfish Cladonema pacificum. PeerJ 7, e7579 (2019).

19. Fujita, S. et al. Distinct stem-like cell populations facilitate functional regeneration of the Cladonema medusa tentacle. PLoS Biol. 21, e3002435 (2023).

20. Kanehisa, R. et al. Aging disrupts tissue homeostasis and constrains blastema-mediated regeneration in the Cladonema medusa. 2026.01.21.700690 Preprint at 10.64898/2026.01.21.700690 (2026).

21. Fujita, S. et al. Division of labor between ERK and Notch signaling coordinates distinct stem cell populations during jellyfish tentacle regeneration. 2026.03.08.710341 Preprint at 10.64898/2026.03.08.710341 (2026).

22. Kondoh, E., Tachibana, K. & Deguchi, R. Intracellular Ca2+ increase induces post-fertilization events via MAP kinase dephosphorylation in eggs of the hydrozoan jellyfish Cladonema pacificum. Dev. Biol. 293, 228–241 (2006).

23. Takeda, N. et al. Identification of jellyfish neuropeptides that act directly as oocyte maturation-inducing hormones. Development 145, dev156786 (2018).

24. Hou, S., Zhu, J., Shibata, S., Nakamoto, A. & Kumano, G. Repetitive accumulation of interstitial cells generates the branched structure of Cladonema medusa tentacles. Development 148, dev199544 (2021).

25. Tachibana, K., Matsumoto, M., Minowa, A. & Deguchi, R. Isolation and Characterization of Feeding-Deficient Strains in Inbred Lines of the Hydrozoan Jellyfish Cladonema pacificum. Zoolog. Sci. 37, 263 (2020).

26. Thoma, V., et al. On the origin of appetite: GLWamide in jellyfish represents an ancestral satiety neuropeptide. Proc. Natl. Acad. Sci. 120, e2221493120 (2023).

27. Mashiba, G., Tanimoto, H. & Thoma, V. Satiety differentially modulates feeding steps in the jellyfish *Cladonema*. iScience 28, 112192 (2025).

28. Takeda, N. et al. Light and Dark Types of Cladonema pacificum (Cnidaria, Hydrozoa): Reproductive Isolation between the Two Types Leading to Speciation. Zoolog. Sci. 43, 236–249 (2026).

29. Marçais, G. & Kingsford, C. A fast, lock-free approach for efficient parallel counting of occurrences of k-mers. Bioinformatics 27, 764–770 (2011).

30. Ranallo-Benavidez, T. R., Jaron, K. S. & Schatz, M. C. GenomeScope 2.0 and Smudgeplot for reference-free profiling of polyploid genomes. Nat. Commun. 11, 1432 (2020).

31. Cheng, H., Concepcion, G. T., Feng, X., Zhang, H. & Li, H. Haplotype-resolved de novo assembly using phased assembly graphs with hifiasm. Nat. Methods 18, 170–175 (2021).

32. Cheng, H. et al. Haplotype-resolved assembly of diploid genomes without parental data. Nat. Biotechnol. 40, 1332–1335 (2022).

33. Cheng, H., Asri, M., Lucas, J., Koren, S. & Li, H. Scalable telomere-to-telomere assembly for diploid and polyploid genomes with double graph. Nat. Methods 21, 967–970 (2024).

34. Zhou, C., McCarthy, S. A. & Durbin, R. YaHS: yet another Hi-C scaffolding tool. Bioinformatics 39, btac808 (2023).

35. Chapman, J. A. et al. The dynamic genome of Hydra. Nature 464, 592–596 (2010).

36. Danecek, P. et al. Twelve years of SAMtools and BCFtools. GigaScience 10, giab008 (2021).

37. Li, H. & Durbin, R. Fast and accurate short read alignment with Burrows-Wheeler transform. Bioinformatics 25, 1754–1760 (2009).

38. Open2C et al. Pairtools: From sequencing data to chromosome contacts. PLoS Comput. Biol. 20, e1012164 (2024).

39. Durand, N. C. et al. Juicer Provides a One-Click System for Analyzing Loop-Resolution Hi-C Experiments. Cell Syst. 3, 95–98 (2016).

40. Durand, N. C. et al. Juicebox Provides a Visualization System for Hi-C Contact Maps with Unlimited Zoom. Cell Syst. 3, 99–101 (2016).

41. Marçais, G. et al. MUMmer4: A fast and versatile genome alignment system. PLOS Comput. Biol. 14, e1005944 (2018).

42. Zacharias, H., Anokhin, B., Khalturin, K. & Bosch, T. C. G. Genome sizes and chromosomes in the basal metazoan *Hydra*. Zoology 107, 219–227 (2004).

43. Simakov, O. et al. Deeply conserved synteny and the evolution of metazoan chromosomes. Sci. Adv. 8, eabi5884 (2022).

44. Tegenfeldt, F. et al. OrthoDB and BUSCO update: annotation of orthologs with wider sampling of genomes. Nucleic Acids Res. 53, D516–D522 (2025).

45. Flynn, J. M. et al. RepeatModeler2 for automated genomic discovery of transposable element families. Proc. Natl. Acad. Sci. 117, 9451–9457 (2020).

46. Tarailo-Graovac, M. & Chen, N. Using RepeatMasker to Identify Repetitive Elements in Genomic Sequences. Curr. Protoc. Bioinforma. 25, 4.10.1–4.10.14 (2009).

47. Hoff, K. J., Lange, S., Lomsadze, A., Borodovsky, M. & Stanke, M. BRAKER1: Unsupervised RNA-Seq-Based Genome Annotation with GeneMark-ET and AUGUSTUS. Bioinformatics 32, 767–769 (2016).

48. Brůna, T., Hoff, K. J., Lomsadze, A., Stanke, M. & Borodovsky, M. BRAKER2: automatic eukaryotic genome annotation with GeneMark-EP+ and AUGUSTUS supported by a protein database. NAR Genomics Bioinforma. 3, lqaa108 (2021).

49. Hoff, K. J., Lomsadze, A., Borodovsky, M. & Stanke, M. Whole-Genome Annotation with BRAKER. Methods Mol. Biol. 1962, 65–95 (2019).

50. Brůna, T., Lomsadze, A. & Borodovsky, M. GeneMark-ETP significantly improves the accuracy of automatic annotation of large eukaryotic genomes. Genome Res. 34, 757–768 (2024).

51. Buchfink, B., Xie, C. & Huson, D. H. Fast and sensitive protein alignment using DIAMOND. Nat. Methods 12, 59–60 (2015).

52. Gotoh, O. A space-efficient and accurate method for mapping and aligning cDNA sequences onto genomic sequence. Nucleic Acids Res. 36, 2630–2638 (2008).

53. Iwata, H. & Gotoh, O. Benchmarking spliced alignment programs including Spaln2, an extended version of Spaln that incorporates additional species-specific features. Nucleic Acids Res. 40, e161 (2012).

54. Kovaka, S. et al. Transcriptome assembly from long-read RNA-seq alignments with StringTie2. Genome Biol. 20, 278 (2019).

55. Pertea, G. & Pertea, M. GFF Utilities: GffRead and GffCompare. F1000Research 9, ISCB Comm J-304 (2020).

56. Gabriel, L. et al. BRAKER3: Fully automated genome annotation using RNA-seq and protein evidence with GeneMark-ETP, AUGUSTUS, and TSEBRA. Genome Res. 34, 769–777 (2024).

57. Stanke, M., Diekhans, M., Baertsch, R. & Haussler, D. Using native and syntenically mapped cDNA alignments to improve de novo gene finding. Bioinformatics 24, 637–644 (2008).

58. Stanke, M., Schöffmann, O., Morgenstern, B. & Waack, S. Gene prediction in eukaryotes with a generalized hidden Markov model that uses hints from external sources. BMC Bioinformatics 7, 62 (2006).

59. Gabriel, L., Hoff, K. J., Brůna, T., Borodovsky, M. & Stanke, M. TSEBRA: transcript selector for BRAKER. BMC Bioinformatics 22, 566 (2021).

60. Dobin, A. et al. STAR: ultrafast universal RNA-seq aligner. Bioinformatics 29, 15–21 (2013).

61. Pacific Biosciences. CLI Workflow. Iso-Seq Docs https://isoseq.how/clustering/cli-workflow.html.

62. Shen, W., Sipos, B. & Zhao, L. SeqKit2: A Swiss army knife for sequence and alignment processing. iMeta 3, e191 (2024).

63. seqtk, Toolkit for processing sequences in FASTA/Q formats. Available from: https://github.com/lh3/seqtk.

64. Camacho, C. et al. BLAST+: architecture and applications. BMC Bioinformatics 10, 421 (2009).

65. UniProt Consortium. UniProt: the Universal Protein Knowledgebase in 2025. Nucleic Acids Res. 53, D609–D617 (2025).

66. Jones, P. et al. InterProScan 5: genome-scale protein function classification. Bioinformatics 30, 1236–1240 (2014).

67. Cantalapiedra, C. P., Hernández-Plaza, A., Letunic, I., Bork, P. & Huerta-Cepas, J. eggNOG-mapper v2: Functional Annotation, Orthology Assignments, and Domain Prediction at the Metagenomic Scale. Mol. Biol. Evol. 38, 5825–5829 (2021).

68. Travert, M. et al. Coevolution of the Tlx homeobox gene with medusa development (Cnidaria: Medusozoa). Commun. Biol. 6, 709 (2023).

69. Katoh, K. & Standley, D. M. MAFFT multiple sequence alignment software version 7: improvements in performance and usability. Mol. Biol. Evol. 30, 772–780 (2013).

70. Capella-Gutiérrez, S., Silla-Martínez, J. M. & Gabaldón, T. trimAl: a tool for automated alignment trimming in large-scale phylogenetic analyses. Bioinformatics 25, 1972–1973 (2009).

71. Wong, T. K. F. et al. IQ-TREE 3: phylogenomic inference software using complex evolutionary models. Mol. Biol. Evol. 43, msag117 (2026).

72. Thorvaldsdóttir, H., Robinson, J. T. & Mesirov, J. P. Integrative Genomics Viewer (IGV): high-performance genomics data visualization and exploration. Brief. Bioinform. 14, 178–192 (2013).

73. Liao, Y., Smyth, G. K. & Shi, W. featureCounts: an efficient general purpose program for assigning sequence reads to genomic features. Bioinformatics 30, 923–930 (2014).

74. Naumov, D. V. Novye rody i vidy gidroidov (Hydroidea) iz more Dal’nego Vostoka. New genera and species of Hydroidea from the Far Eastern Seas. Tr. Zool. Instituta Akad. Nauk SSSR Proc. Zool. Inst. USSR Acad. Sci. 18: 19–25 (1955).

75. Emms, D. M. & Kelly, S. OrthoFinder: phylogenetic orthology inference for comparative genomics. Genome Biol. 20, 238 (2019).

76. Castresana, J. Selection of conserved blocks from multiple alignments for their use in phylogenetic analysis. Mol. Biol. Evol. 17, 540–552 (2000).

77. Talavera, G. & Castresana, J. Improvement of Phylogenies after Removing Divergent and Ambiguously Aligned Blocks from Protein Sequence Alignments. Syst. Biol. 56, 564–577 (2007).

78. Borowiec, M. L. AMAS: a fast tool for alignment manipulation and computing of summary statistics. PeerJ 4, e1660 (2016).

79. Kumar, S. et al. MEGA12: Molecular Evolutionary Genetic Analysis Version 12 for Adaptive and Green Computing. Mol. Biol. Evol. 41, msae263 (2024).

80. Hilali, S. E. & Copley, R. R. macrosyntR : Drawing automatically ordered Oxford Grids from standard genomic files in R. 2023.01.26.525673 Preprint at 10.1101/2023.01.26.525673 (2023).

81. DNA Data Bank of Japan. https://ddbj.nig.ac.jp/search/entry/bioproject/PRJDB42964 (2026).

82. DNA Data Bank of Japan. https://ddbj.nig.ac.jp/search/entry/bioproject/PRJDB45818 (2026).

83. DNA Data Bank of Japan. https://ddbj.nig.ac.jp/search/entry/bioproject/PRJDB42990 (2026).

84. DNA Data Bank of Japan. https://ddbj.nig.ac.jp/search/entry/bioproject/PRJDB42987 (2026).

85. DNA Data Bank of Japan. https://ddbj.nig.ac.jp/search/entry/bioproject/PRJDB42972 (2026).

86. DNA Data Bank of Japan. https://ddbj.nig.ac.jp/search/entry/bioproject/PRJDB42991 (2026).

87. DNA Data Bank of Japan. https://identifiers.org/ncbi/insdc:BAANGU000000000 (2026).

88. Ui, J., et al. Chromosome-level genome assembly of the hydrozoan jellyfish Cladonema pacificum. figshare 10.6084/m9.figshare.35116864.v1 (2026).

89. Chen, S., Zhou, Y., Chen, Y. & Gu, J. fastp: an ultra-fast all-in-one FASTQ preprocessor. Bioinformatics 34, i884–i890 (2018).

